# Contribution of dorsal versus ventral hippocampus to the hierarchical modulation of goal-directed action

**DOI:** 10.1101/2023.05.23.541867

**Authors:** Robin Piquet, Angélique Faugère, Shauna L. Parkes

## Abstract

Adaptive behavior often necessitates that animals learn about events in a manner that is specific to a particular context or environment. These hierarchical organizations allow the animal to decide which action is the most appropriate when faced with ambiguous or conflicting possibilities. This study examined the role of hippocampus in enabling animals to use the context to guide action selection. We used a hierarchical instrumental outcome devaluation task in which male rats learn that the context provides information about the unique action-outcome relations that are in effect. We first confirmed that rats encode and use hierarchical context-(action-outcome) relations. We then show that chemogenetic inhibition of ventral hippocampus (vHPC) impairs both the encoding and retrieval of these associations, while inhibition of dorsal hippocampus (dHPC) impairs only the retrieval. Importantly, neither dHPC or vHPC were required for goal-directed behavior *per se* as these impairments only emerged when rats were forced to use the context to identify the current action-outcome relationships. These findings are discussed with respect to the role of the hippocampus and its broader circuitry in the contextual modulation of goal-directed behavior and the importance of hierarchical associations in flexible behavior.

## Introduction

Animals readily encode associations between actions and outcomes, but these simple binary associations do not fully explain the complexity of action selection (Rescorla, 1992). Indeed, animals can also learn hierarchical organizations whereby a context, or stimulus, comes to modulate the action-outcome association (Colwill and Rescorla, 1990; Rescorla, 1992; Haddon and Killcross, 2006a, 2007; Marquis et al., 2007; Haddon et al., 2008; Bradfield and Balleine, 2013; Trask and Bouton, 2014) and to function as a “gatekeeper” to select the most appropriate behavioral action (Bouton and Swartzentruber, 1986; Myers and Gluck, 1994; Lee and Lee, 2013)

Colwill and Rescorla (1990) first provided evidence for hierarchical associations by training rats in a task in which two discrete stimuli (S) dictated that distinct action-outcome (A-O) associations would be in effect. For example, rats learned that, in the presence of S1, A1 earned O1 and A2 earned O2, but these relations were reversed in the presence of S2 (S2: A1-O2; A2-O1). Following devaluation of O1, rats selectively reduced their responding on A1 when S1 was present and on A2 when S2 was present. Later, Trask and Bouton (2014) showed that the context can enter into a similar hierarchical association and signal that an action will produce a specific outcome in a specific context. Rats were trained in one context (Context A) in which responding on A1 earned O1 and responding on A2 earned O2 and in a different context (Context B) where responding on A1 earned O2 and A2 earned O1. At test, after devaluation of O1, rats reduced performance on A1 in Context A and A2 in Context B. This indicates that rats had learnt specific A-O associations within a particular context, and suggests that animals can learn and use context-(A-O) hierarchical relationships.

Together, these studies show that hierarchical associations are pervasive in learning and are used to guide goal-directed actions when simple binary A-O associations do not suffice (Thrailkill and Bouton, 2015). Importantly, the neural bases of context-basedhierarchical associations remain largely unexplored. One clear candidate is the hippocampus as it has long been implicated in the contextual modulation of behavior (Rudy, 2009; Maren et al., 2013) and, in parallel, its role in goal-directed behavior is receiving renewed attention (Corbit et al., 2002; Gourley et al., 2010; Barfield et al., 2017; Barfield and Gourley, 2019; Yoshida et al., 2019, 2021; Bradfield et al., 2020). The dichotomy between the vHPC and dHPC in the contextual regulation of behavior has also been the focus of several studies (e.g., Wang et al., 2013; Riaz et al., 2017; Park et al., 2020; Pinizzotto et al., 2020; Oleksiak et al., 2021; Biane et al., 2023). For example, inhibition of the ventral hippocampus (vHPC), but not dorsal hippocampus (dHPC), prevents rats to respond to the appropriate cue in the appropriate context in a Pavlovian biconditional discrimination task (Riaz et al., 2017) and impairs context-dependent active avoidance (Oleksiak et al., 2021).

We therefore examined the possible role of vHPC and dHPC in the formation and retrieval of hierarchical context-(action-outcome) associations. We confirm and extend previous findings that context can modulate action-outcome associations and be used to guide action selection (Trask and Bouton, 2014). Using chemogenetic inhibition, we also demonstrate that the vHPC is required to both encode and retrieve this hierarchical association, whereas dHPC is required only for retrieval. Importantly, we show that goal-directed behavior per se is not impacted by chemogenetic inhibition of vHPC or dHPC but, instead, these regions appear to play a specific role in the contextual regulation of goal-directed behavior.

## Materials and Methods

### Subjects

Subjects were 79 male Long-Evans rats aged 3-4 months (Janvier, France). The rats used in Experiment 1a and 2a (n = 16) had been previously trained in an unrelated instrumental conditioning task but all other rats were naïve. Rats were housed in pairs in plastic boxes located in a climate controlled room maintained on a 12 h light/dark cycle (lights on at 07:00). Behavior occurred during the light phase of the cycle. Rats were handled daily for five days before the behavioral procedures and put on food restriction two days before behavior to maintain them at approximately 90% of their *ad libitum* feeding weight. Experiments were conducted in agreement with French (council directive 2013-118, February 1, 2013) and European (directive 2010-63, September 22, 2010, European Community) legislations and received approval from the local Ethics Committee (CE50).

### Viral vectors

In Experiments 1b and 2b, an adeno-associated viral vector carrying the inhibitory hM4Di designer receptor exclusively activated by designer drugs (DREADD; Armbruster et al., 2007; Rogan and Roth, 2011) was obtained from Viral Vector Production Unit (Universitat Autonoma de Barcelona, Spain) using a plasmid obtained from Addgene (pAAV-CaMKIIa-hM4D(Gi)-mCherry; Addgene plasmid # 50477; http://n2t.net/addgene:50477; RRID:Addgene_50477; gift from Bryan Roth). The vector used was AAV8-CaMKII-hM4D(Gi)-mCherry (1.46 x 10^13^ gc/ml). A control vector lacking the hM4Di receptor was also used (AAV8-CaMKII-EGFP; 2.1 x 10^13^ gc/ml Addgene viral prep # 50469-AAV8; http://n2t.net/addgene:50469; RRID:Addgene_50469; plasmid was a gift from Bryan Roth). The DREADD ligand, deschloroclozapine (DCZ; HY-42110, MedChemExpress) was dissolved in dimethyl sulfoxide (DMSO) to a final volume of 50 mg/ml and stored at -80°C. This solution was then diluted (1:500) in sterile saline to a final concentration of 0.1 mg/ml and was injected intraperitoneally (i.p.) 30 min before behavior. New solutions were prepared each day. DCZ was prepared and injected under low light conditions. We, and others, have previously demonstrated the efficacy of this ligand (Nagai et al., 2020; Nentwig et al., 2021; Cerpa et al., 2023). The vehicle solution was composed of 0.2% DMSO in sterile saline.

### Surgery

Rats were anaesthetized using Isoflurane (5% induction; 1-2% maintenance) and mounted on a stereotaxic apparatus (Kopf). The incision site was subcutaneously injected with 0.2 ml of local anaesthetic (ropivicaine) and then disinfected using betadine. The viral vectors were injected using a 10 µl Hamilton syringe connected to a microinjector (UMP3 UltraMicroPump II with Micro4 Controller, World Precision Instruments). 0.8 µl of AAV was injected at a rate of 0.2 µl/min at two sites in each hemisphere, i.e., 1.6 µl per hemisphere. The co-ordinates for the ventral hippocampus were: AP -5.4, ML ±5.3, DV -6.5, and AP - 6.0, ML ±4.6, DV -8.2 (mm from bregma; Paxinos and Watson, 2013) and the coordinates for dorsal hippocampus were: AP -3.5, ML ±1.4, DV -3.0, and AP -3.8, ML ±3.0, DV -2.5. For each experiment, half of the rats were injected with AAV8-CaMKII-hM4D(Gi)-mCherry (hM4Di groups) and the other half were injected with the control virus AAV8-CaMKII-EGFP (GFP groups). During surgery, a heating pad was placed under the rat to maintain body temperature and rats were rehydrated with subcutaneous injections of warm saline (0.9%, 10 ml/kg/hour). At the end of surgery, rats were subcutaneously injected with a nonsteroidal anti-inflammatory drug (meloxicam, 2mg/ml/kg) and were individually housed in a warm cage with facilitated access to food and water for 1-2 h post-surgery. All rats were given at least 5 days of post-operative care and were allowed a minimum of 4 weeks to recover before the start of the behavioral procedures.

### Behavioral Apparatus

Instrumental training and testing took place in two sets of 8 operant cages (40 cm width x 30 cm depth x 35 cm height, Imetronic, France) located in different experimental rooms. The first set of cages (Context A) had grey (sides) and transparent (front and back) plastic walls. The floor was made of stainless steel rods and a plastic divider with a rounded edge (12 cm length x 25 cm height) was added to each cage. A 10% lemon scent was added to the bedding below the floor before each session (Arôme citron, Vahiné, France) and the inside of the cages was cleaned with water between each rat. The second set of cages (Context B) were located in a different room of the laboratory. The walls were grey (sides) and transparent (front and back) with a vertical black and white striped pattern added to the back wall and a black and white diamond pattern added to the front wall. The floor was smooth solid plastic with a black and white checkered pattern and two led lights were added to the top of one side wall.

All operant cages were equipped with two pellet dispensers that delivered grain or sugar pellets into a single food port when activated. The cages also contained two retractable levers that could be inserted to the left and right of the food port, and a house light illuminated the cage. Experimental events were controlled and recorded by a computer located in the room. Outcome devaluation occurred in individual polycarbonate feeding cages located in a different room to the operant cages.

### Behavioral Procedures

The behavioral procedure used in all experiments is illustrated in Figure 1A and is described in detail below.

**Figure 1.**
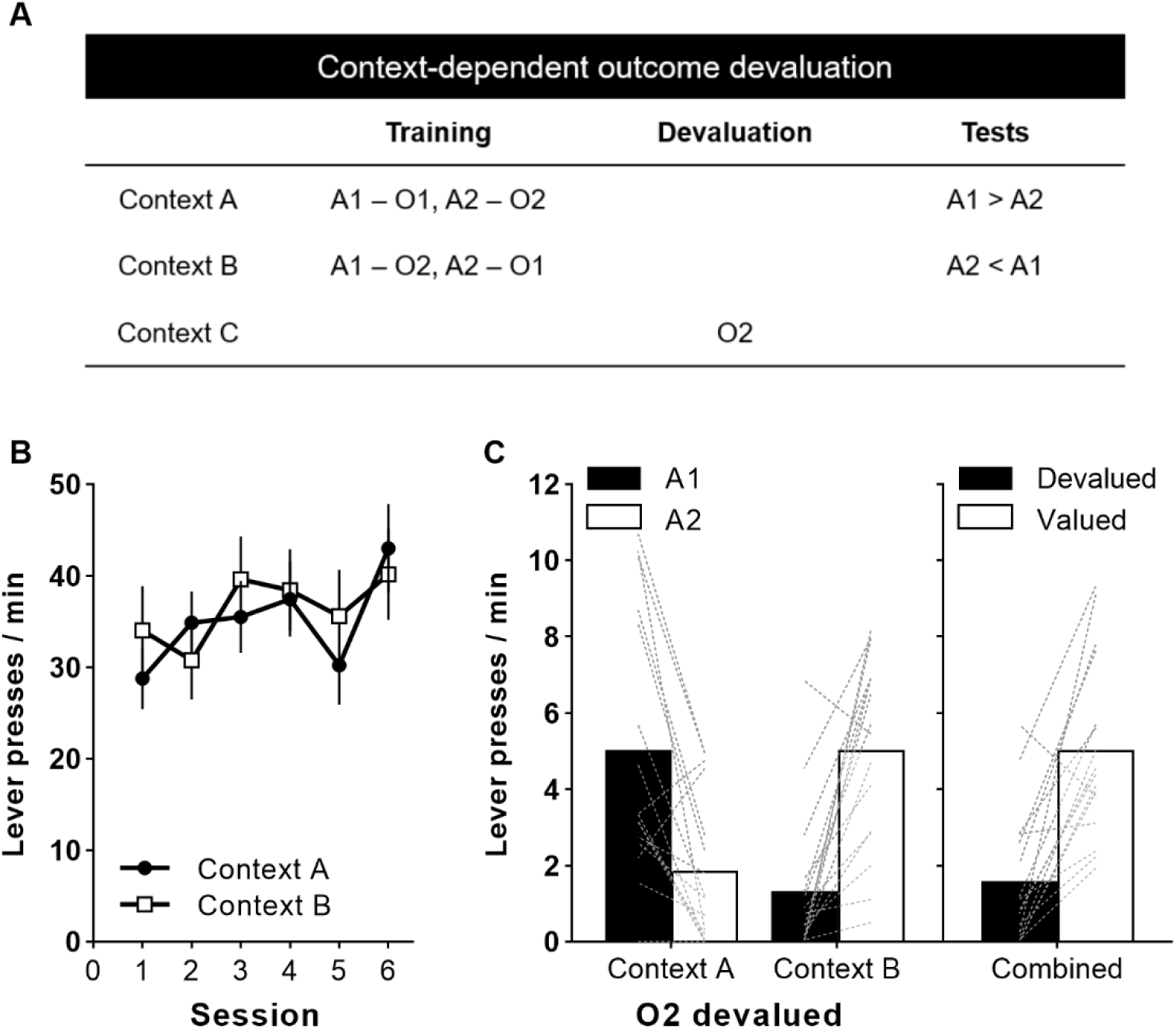
**(A)** Schematic of the behavioral design that was used in all experiments (Trask & Bouton, 2014). A1 and A2 indicate counterbalanced left or right levers and O1 and O2 represent counterbalanced grain or sugar pellets. Context A and B are different sets of operant chambers that have distinct contextual cues. **(B)** Rate of lever press responding (±SEM) during instrumental training in Context A and B. Performance is averaged across the two levers. **(C)** Average (±SEM) lever presses per minute on A1 and A2 in each context (left panel) and collapsed across contexts according to whether the action was associated with the devalued or valued outcome (right panel).

#### Instrumental training

On the first three days, rats were trained to enter the recessed food port to retrieve grain and sugar pellets that were delivered every 60 s on average. Rats had two daily sessions, one each context with the order counterbalanced. The session duration was 20 min during which they received 10 grain and 10 sugar pellets in each context. Levers were retracted during these sessions. Over the next 9 days, rats were given two instrumental training sessions per day, one in each context with the order counterbalanced. These sessions were separated by 10 min. In each context, rats were trained to perform one lever (e.g., left lever) to earn one outcome (e.g., grain pellet) and the other lever (e.g., right lever) to earn the other outcome (e.g. sugar pellet). Importantly, these action-outcome associations were reversed in each context. For example, a rat trained in Context A to press the left lever for grain and the right lever for sugar would also be trained in Context B to press the left lever for sugar and the right lever for grain. The action-outcome associations were fully counterbalanced across rats and contexts.

Each session lasted 25 min maximum, during which each lever was presented for 10 min or until 20 pellets had been earned. Each session began with a 2 min acclimation period to the context before the first lever was presented. The interval between each lever presentation was 2 min and rats remained in the cage for 1 min once the second lever had retracted. The order of context presentation was counterbalanced between rats and across days. The order of lever presentation across days was counterbalanced in a pseudo-randomized sequence. The same lever was presented first in Context A and B on any given day. For the first 3 days, rats were trained under continuous reinforcement (CRF), then under an RR5 schedule (on average, rewards were delivered after 5 presses) for 3 days, and under an RR10 schedule (on average, reward delivered after 10 presses) for the last 3 days.

As stated above, the rats used in Experiment 1a and 2a had previously participated in an unrelated instrumental task. These rats therefore received only 6 sessions of instrumental training in each context (12 sessions total) rather than 9 sessions per context (18 sessions total). In chemogenetic experiments, half of the rats in each group received an i.p. injection of vehicle and the other half received an injection of the DREADD ligand, DCZ, 30 min prior to the first instrumental training session each day.

#### Outcome devaluation

Twenty-four hours after the final instrumental training session, rats underwent outcome-specific devaluation via sensory specific satiety during which they were given 1 h access to one of the outcomes (identity of the devalued outcome was fully counterbalanced within groups) in plastic feeding cages located in a different room to the operant cages. After devaluation, rats were given two consecutive choice tests (one in each context). The interval between each test was 10 min and the order of the tests was fully counterbalanced. After a 2 min acclimation period, both levers were presented simultaneously for 10 min and responding on the levers was unrewarded. Immediately after the second test, rats were given a consumption preference test to assess the effectiveness of the satiety-induced devaluation. Rats were placed back in the plastic feeding cages and were given 10 min access to both food outcomes. Forty-eight hours later, rats received a second series of tests with the other outcome devalued.

In chemogenetic experiments, rats that received DCZ during training received vehicle at test and rats that received vehicle during training received an injection of DCZ at test. The injections were performed after the satiety session and rats were placed in the operant cages for the unrewarded tests 30 min after injection.

### Immunofluorescence for mCherry

Subsequent to behavioral testing, rats were rapidly and deeply anaesthetized with pentobarbital monosodic and perfused transcardially with 4% paraformaldehyde in 0.1M phosphate buffer. Brains were removed and post-fixed in 4% paraformaldehyde overnight. Subsequently, 40 µm coronal sections were cut using a VT1200S Vibratome (Leica Microsystems). Every fourth section was collected to form a series and immunoreactivity was performed for mCherry. Free-floating sections were prepared by rinsing in 0.1M phosphate buffered saline with 0.3% Triton X-100 (PBST) for 4*5 min, incubated in 0.5% H_2_O_2_ diluted in 0.1M PBST, rinsed in 0.1M PBST for 4*5 min, blocked (1 h, PBST 0.1M, 4% normal goat serum) and placed in 1:1000 rabbit anti-RFP (red fluorescent protein; PM005 CliniSciences; RRID:AB_591279) diluted in 0.1M PBST at room temperature overnight. Sections were then rinsed, incubated in 1:500 Biotin-SP AffiniPure Goat Anti-Rabbit IgG (Jackson Immunoresearch; 111-065-003; RRID: AB_2337959) diluted in 0.1M PBST for 2 h at room temperature, rinsed again, and then placed in 1:400 Alexa Fluor® 594 Streptavidin (Jackson Immunoresearch; 016-580-084; RRID: AB_2337250) diluted in 0.1M PBS. Sections were then rinsed and incubated in 1:5000 bisBenzimide H 33258 (Sigma-Aldrich; 14530) diluted in 0.1M PBS for 15 min. Finally, sections were rinsed, mounted, and cover-slipped with Fluoromount-G (SouthernBiotech). Sections were then imaged using a Nanozoomer slide scanner (Hamamatsu Photonics) and analyzed with the NDP.view 2^®^ freeware (Hamamatsu Photonics).

### Statistical Analyses

Experiments 1a and 2a used a 2 x 2 within-subjects design (devalued versus valued and Context A versus Context B). Experiments 1b and 2b used a mixed methods design with two between-subject factors (group: hM4Di or EGFP; treatment: DCZ or vehicle) and a within-subject factor of devaluation (devalued versus non-devalued). Data were analysed using sets of between and within orthogonal contrasts controlling the per contrast error rate at alpha = 0.05 (Harris, 1994; Hays, 1963). Simple effects analyses were conducted to establish the source of significant interactions. Statistical analyses were performed using PSY Statistical Program (http://www.psy.unsw.edu.au/research/research-tools/psy-statistical-program; Kevin Bird, Dusan Hadzi-Pavlovic, and Andrew Issac © School of Psychology, University of New South Wales) and graphs were created using GraphPad

Prism. Statistical significance was set at p ≤ 0.05. Data are presented as mean ± SEM.

## Results

### Experiment 1a

We first aimed to replicate the findings of Trask and Bouton (2014) using the experimental design shown in Figure 1A. Briefly, rats were trained in different contexts (Context A and Context B) to perform two different actions (A1 and A2) to earn two distinct food outcomes (O1 and O2). Importantly, the action-outcome (A-O) associations were reversed in each context. One of the outcomes was then devalued via sensory specific satiety in a third context (Context C) and, finally, rats were tested in both Context A and Context B, with the order counterbalanced. It was predicted that rats would show differential responding in the two contexts. For example, if O2 was devalued, rats should press more on A1 in Context A but more on A2 in Context B.

Instrumental training occurred without incident and the rate of lever pressing in the two contexts did not differ (F_1,15_ = 0.33; p = 0.57; Figure 1B). The test results are shown in Figure 1C. Statistics revealed no within-subjects effect of context (F_1,15_ = 0.2; p = 0.66) or lever (F_1,15_ = 0.42; p = 0.53) but there was a significant interaction between these factors (F_1,15_ = 30.96; p < 0.001), indicating that responding on each lever depended on the test context. The overall analysis of the combined data (right panel), collapsed across action and context, showed that rats pressed significantly less on the lever that was associated with the devalued outcome (F_1,15_ = 31.0; p < 0.001). The rats also consumed significantly less of the devalued outcome compared to the non-devalued outcome during the subsequent consumption test (F_1,15_ = 103.45; p < 0.001; data not shown). These results replicate Trask and Bouton (2014) and show that rats can select the action associated with the still valued outcome according to the context.

### Experiment 1b

The next experiment assessed the impact of chemogenetic inhibition of ventral hippocampus (vHPC) or dorsal hippocampus (dHPC) using the experimental design shown in Figure 1A. Rats were injected i.p. with DCZ or vehicle during training or during the tests.

#### Histology

A representative image of viral expression in vHPC is shown in Figure 2A and a schematic of the viral expression for each rat is illustrated in Figure 2B. One rat was excluded due to misplaced injections (injections were too dorsal and did not reach the ventral tip of CA1 and ventral subiculum). This yielded the following between-subject group sizes: DCZ at training: GFP n = 8, hM4Di n = 7, and DCZ at test: GFP *n* = 8, hM4Di n = 8. Figure 2C shows a representative image of viral expression in dHPC and the extent of viral expression for all rats is shown in Figure 2D. One rat was excluded due to misplaced injections (injections were too dorsal leading to almost no infected cells in dHPC but extensive expression in the cortex), yielding the following between-subject group sizes: DCZ at training: GFP n = 8, hM4Di n = 7, and DCZ at test: GFP *n* = 8, hM4Di n = 7.

**Figure 2.**
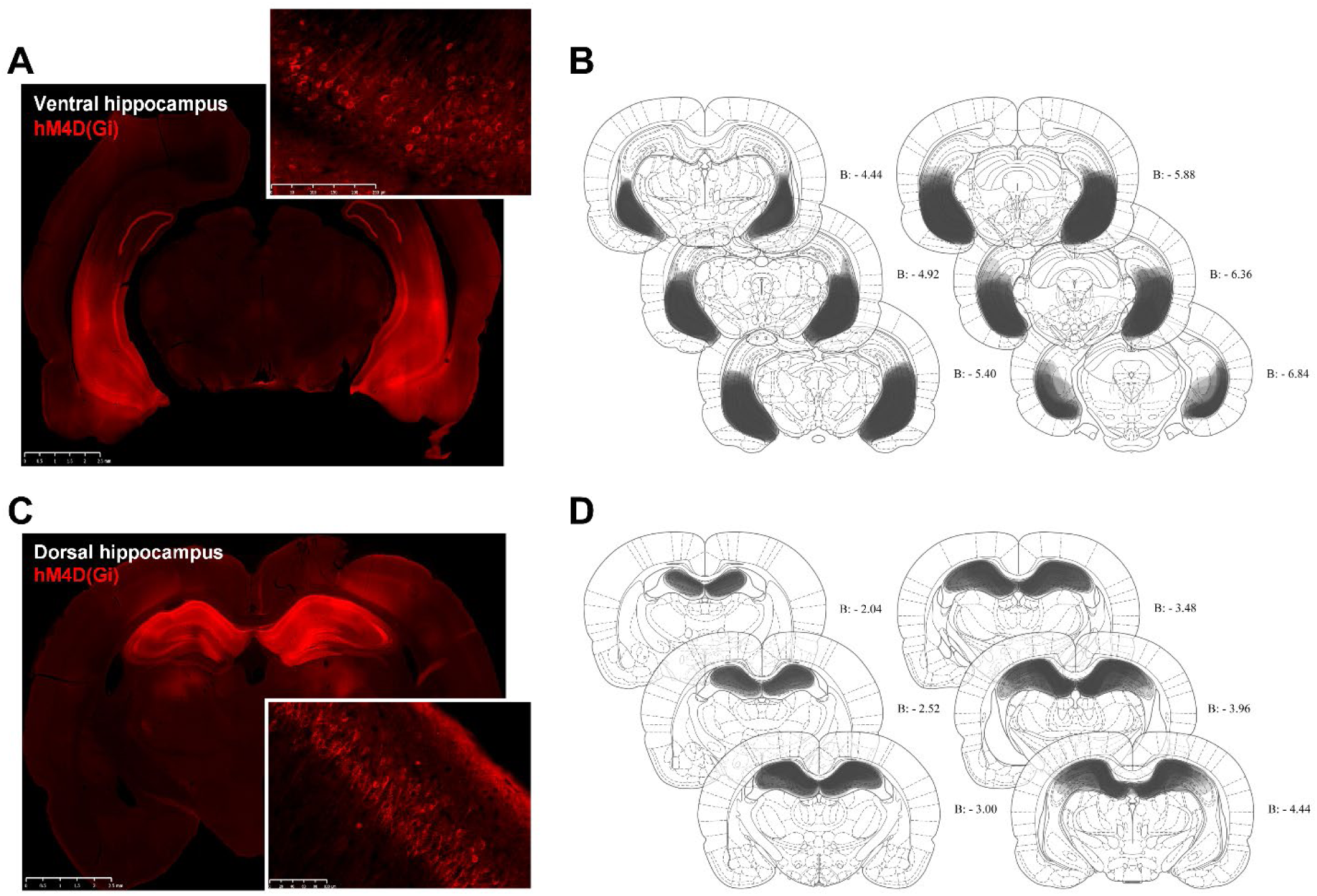
**(A, C)** Representative photomicrographs of the expression of hM4D(Gi)-mCherry in ventral and dorsal hippocampus, respectively. **(B, D)** Illustrations of the mCherry expression for all rats injected in ventral and dorsal hippocampus, respectively. Shading indicates the extent of virus expression, with each rat represented as a separate, stacked layer.

#### Ventral hippocampus

Figure 3A and 3B show the results from rats that received viral injections in vHPC. Rats increased their rate of lever pressing across instrumental training (F_1,27_ = 338.69; p < 0.001; Figure 3A) and there was no effect of virus (hM4Di vs GFP; F_1,27_ = 0.33; p = 0.57) or treatment (DCZ vs vehicle; F_1,27_ = 0.03; p = 0.86) and no significant interaction between these factors (largest F_1,27_ = 1.94; p = 0.18). The test data (Figure 3B) is presented collapsed across contexts as responding did not significantly differ between contexts (context x devaluation interaction: F_1,27_ = 3.19; p = 0.09). Inspection of the figure suggests that all groups showed goal-directed behavior and reduced their responding on the lever associated with the devalued outcome. Statistical analyses confirmed an overall within-subject effect of devaluation (F_1,27_ = 194.58; p < 0.001), and no effect of virus (F_1,27_ = 1.24; p = 0.28) or treatment (F_1,27_ = 1.11; p = 0.30). There was a significant devaluation x treatment interaction (F_1,27_ = 13.66; p = 0.001) but simple effect analyses confirmed a significant devaluation effect in all 4 groups (DCZ at training: GFP, F_1,27_ = 90.08; p < 0.001; hM4Di, F_1,27_ = 62.71; p < 0.001; and DCZ at test: GFP, F_1,27_ = 45.58; p < 0.001; hM4Di, F_1,27_ = 13.569; p = 0.001). During consumption tests, rats consumed significantly less of the devalued outcome compared to the non-devalued outcome (F_1,27_ = 53.17; p < 0.001; data not shown) and there was no effect of virus (F_1,27_ = 0.11; p = 0.74) or treatment (F_1,27_ = 1.16; p = 0.29). No significant interactions between these factors were detected (largest F_1,27_ = 0.69; p = 0.41).

**Figure 3.**
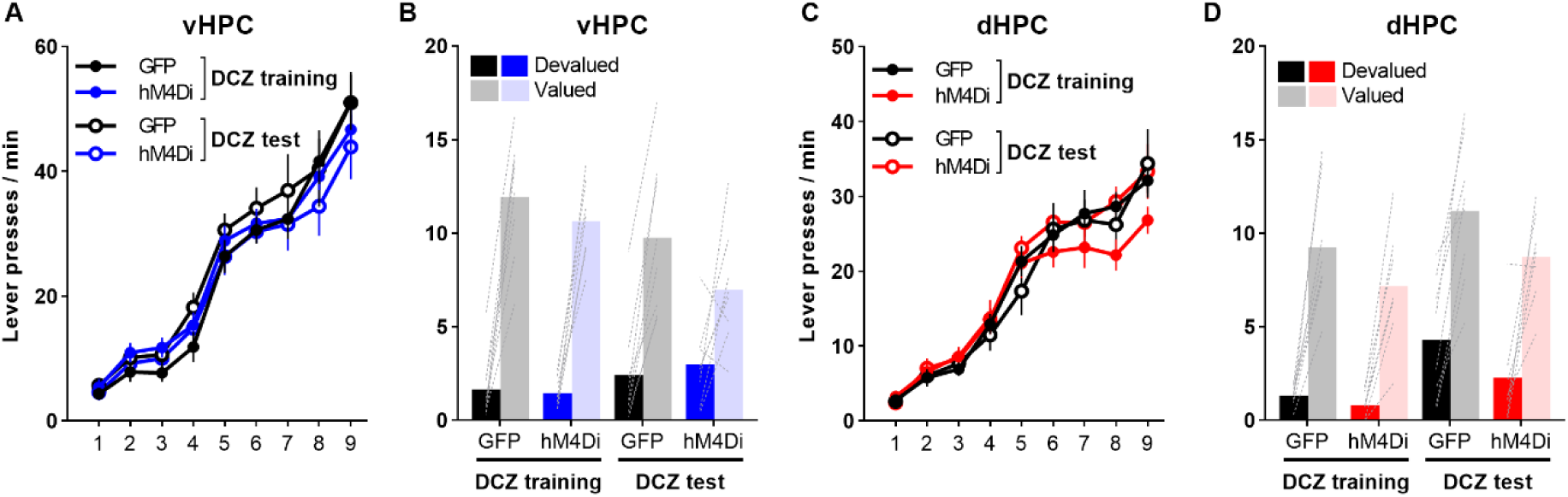
**(A, C)** Average rate of lever pressing (±SEM) across instrumental training averaged across the two levers and the two contexts for rats injected with virus in the ventral hippocampus (vHPC) and dorsal hippopcampus (dHPC), respectively. Rats were injected with either vehicle (open circles) or DCZ (closed circles) before each training session. **(B, D)** Average lever presses per minute (±SEM) during the outcome devaluation test, averaged across the two contexts. Rats were injected with either vehicle or the DREADD ligand, DCZ, before each test.

#### Dorsal hippocampus

Figure 3C and 3D show the results from rats that received the viral injections in dHPC. These rats also increased their performance across instrumental training sessions (F_1,26_ = 352.14; p < 0.001; Figure 3C) and there was no effect of virus (hM4Di vs GFP; F_1,26_ = 0.008; p = 0.93), treatment (DCZ vs vehicle; F_1,26_ = 0.28; p = 0.60), or any interactions between these factors (largest F_1,26_ = 1.56; p = 0.22). Test data is shown in Figure 3D and is presented collapsed across contexts, as responding did not significantly differ between contexts (context x devaluation interaction: F_1,26_ = 1.90; p = 0.18). It appears that all groups pressed less on the lever associated with the devalued outcome compared to the lever associated with the still valued outcome. Statistical analyses confirmed this observation and revealed an overall within-subject effect of devaluation (F_1,26_ = 140.10; p < 0.001). There was also a significant main effect of virus (F_1,26_ = 4.36; p = 0.047) and treatment (F_1,26_ = 5.63; p = 0.025) but no significant interactions (largest F_1,26_ = 0.75; p = 0.39). This indicates that hM4Di groups responded less than GFP groups (regardless of treatment) and rats given vehicle responded less than rats given DCZ (regardless of virus type). Rats in all groups also consumed significantly less of the devalued outcome compared to the non-devalued outcome during the subsequent consumption tests (F_1,26_ = 65.95; p < 0.001; data not shown) and there was no effect of virus (F_1,26_ = 0.21; p = 0.65), or treatment (F_1,26_ = 0.96; p = 0.34), or any interaction between these factors (largest F_1,26_ = 2.14; p = 0.16).

Together, these results suggest that inhibition of ventral or dorsal hippocampus had no effect on the rats’ ability to perform in a context-dependent manner. These results were surprising, however, they should be interpreted with caution. Indeed, in the current experiment, and that of Trask and Bouton (2014), Context A and B were two distinct sets of operant cages. This means that the contexts differed not only in their configural information (olfactory, visual, and tactile) but also in any discrete cues associated to the specific actions used in each context. Rats may therefore have used these cues to guide their responding in each context or to form distinct sets of binary A-O associations in each context (i.e., A1-O1, A2-O2 in context A and A3-O1, A4-O2 in context B) (Bradfield and Balleine, 2013).

In the next experiments, we therefore attempted to dissociate the influence of context versus action-specific cues on responding. To do this, we wanted to remove any action-specific cues during the test session while still preserving the contextual information. We therefore tested the same rats in a new operant cage that contained the relevant contextual information but unfamiliar levers. We hypothesized that, when tested in these cages, rats cannot use any action-specific cues from training to signal the A-O associations in effect and, thus, would be forced to rely on hierarchical context-(A-O) associations formed during training to guide their choice.

### Experiment 2a

We first behaviorally confirmed that the contextual control of instrumental behavior can be transferred to new environments and that hierarchical context-(A-O) associations are maintained even in the absence of action-specific cues. Rats from Experiment 1a were retrained on an RR10 schedule for two days and then underwent a second series of tests in new cages that contained the relevant contextual cues but unfamiliar levers. Figure 4A shows the results from this test. When split by context (left panel), there was no main effect of context (F_1,15_ < 0.001; p > 0.98) or lever (F_1,15_ = 0.002; p = 0.97) but a significant context by lever interaction (F_1,15_ = 17.65; p = 0.001) indicating that responding on each lever differed depending on the context. The combined data from both contexts (right panel) confirmed that rats pressed less on the lever associated with the devalued outcome (F_1,15_ = 17.65; p = 0.001) and the rats also consumed less of the devalued outcome during the consumption test (F_1,15_ = 53.28; p < 0.001; data not shown). These results demonstrate that the context itself can indeed enter into hierarchical relationships with instrumental actions and can be used to signal the specific A-O that are in effect, even in the absence of action-specific cues.

**Figure 4.**
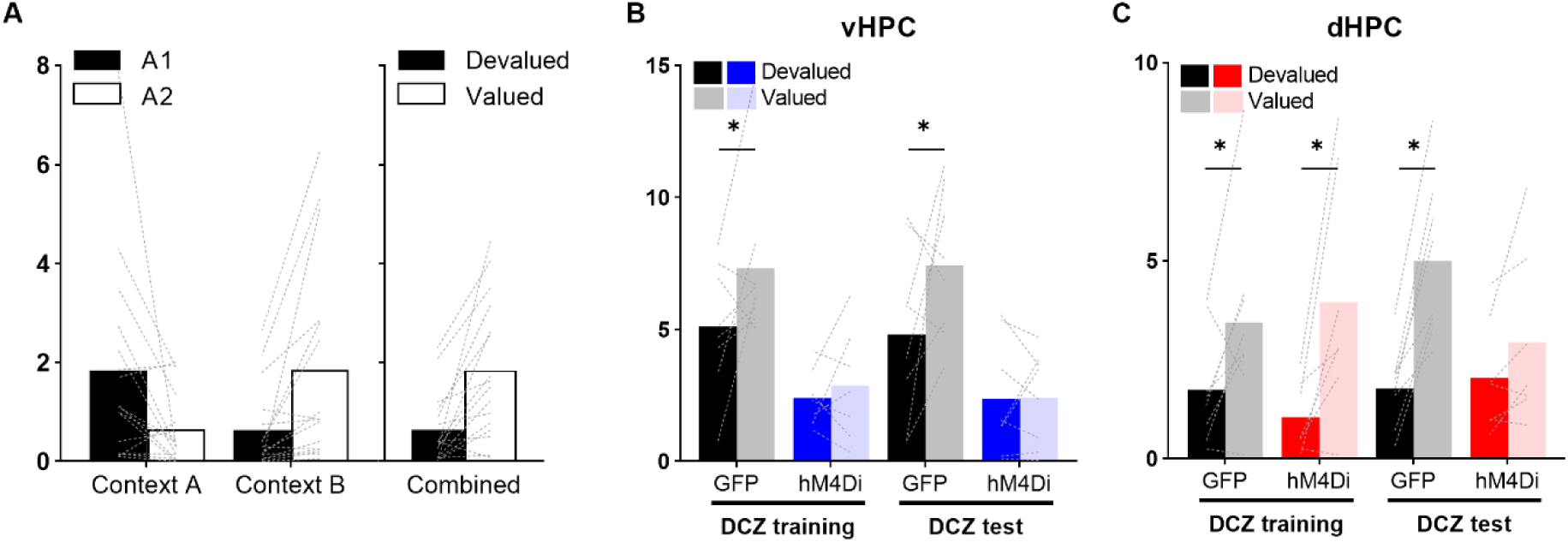
**(A)** Average (±SEM) lever presses per minute on A1 and A2 in each context (left panel) and collapsed across contexts according to whether the action was associated with the devalued or valued outcome (right panel). **(B, C)** Average lever presses per minute (±SEM) during the outcome devaluation test, averaged across the two contexts, for rats with virus in ventral hippocampus (vHPC) or dorsal hippocampus (dHPC), respectively. Rats were injected with either vehicle or DCZ before each test.

### Experiment 2b

We next retrained the rats from Experiment 1b on an RR10 schedule for two days and then tested them in the new cages that contained the relevant contextual cues. Rats were injected with either DCZ or vehicle before each retraining and test session.

#### Ventral hippocampus

Figure 4B shows the results from rats that received viral injections in vHPC. Data are presented averaged across contexts as there was no context x devaluation interaction (F_1,27_ = 0.14; p = 0.71). Statistical analyses revealed an overall within-subject effect of devaluation (F_1,27_ = 6.62; p = 0.02) and virus (F_1,27_ = 25.71; p < 0.001) but no effect of treatment (F_1,27_ = 0.06; p = 0.81). Importantly, there was also a significant devaluation x virus interaction (F_1,27_ = 4.27; p = 0.049). Simple effect analyses revealed a devaluation effect in both GFP groups (DCZ at training: F_1,27_ = 4.69; p = 0.039; DCZ at test: F_1,27_ = 6.53; p = 0.017) but not in the hM4Di groups (DCZ at training: F_1,27_ = 0.19; p = 0.67; DCZ at test: F_1,27_ = 0.003; p = 0.96). That is, selective outcome devaluation was impaired when vHPC was inhibited during training or during test. The satiety-induced devaluation was effective for all groups as, during consumption tests, all rats consumed significantly less of the devalued outcome compared to the non-devalued outcome (F_1,27_ = 245.80; p < 0.001; data not shown), with no significant effect of virus (F_1,27_ = 0.81; p = 0.38) or treatment (F_1,27_ = 2.61; p = 0.12), or any interactions between these factors (largest F_1,27_ = 0.35; p = 0.56). These results indicate that vHPC is required for both the encoding and retrieval of context-(A-O) associations.

#### Dorsal hippocampus

Figure 4C shows the results from rats that received the viral injections in dHPC. Again, data are presented collapsed across contexts as the context x devaluation interaction was not significant (F_1,26_ = 1.90; p = 0.18). Inspection of the figure suggests that inhibition of dHPC during test, but not during training, rendered rats unable to bias their responding toward the appropriate lever. Statistical analyses confirmed this observation and revealed an overall within-subject effect of devaluation (F_1,26_ = 41.88; p < 0.001), no effect of virus (F_1,26_ = 0.63; p = 0.44) or treatment (F_1,26_ = 0.40; p = 0.53), but a significant three-way devaluation x virus x treatment interaction (F_1,26_ = 6.85; p = 0.015). Simple effect analyses revealed a devaluation effect in both GFP groups (DCZ at training: F_1,26_ = 6.80; p = 0.015; DCZ at test: F_1,26_ = 24.48; p < 0.001) and in the hM4Di group that received DCZ at training and vehicle at test (F_1,26_ = 17.41; p < 0.001) but not in the hM4Di group that received DCZ at test (F_1,26_ = 1.67; p = 0.21). All rats consumed significantly less of the devalued outcome compared to the non-devalued outcome during the subsequent consumption tests (F_1,26_ = 100.83; p < 0.001; data not shown), with no effect of virus (F_1,26_ = 0.33; p = 0.57), treatment (F_1,26_ = 2.88; p = 0.10), or any significant interactions (largest F_1,26_ = 0.63; p = 0.44). This finding demonstrates that the dHPC is specifically required for the retrieval of context-(A-O) associations as its inhibition at test, but not during training, impaired context-dependent outcome devaluation.

## Discussion

The results of the current study show that hippocampus plays a central role in the contextual modulation of goal-directed behavior. Our data confirm that rats encode and use hierarchical context-(action-outcome) associations and show that inhibition of ventral hippocampus (vHPC) impairs both the formation and retrieval of these associations, but inhibition of dorsal hippocampus (dHPC) impairs only the retrieval. Importantly, chemogenetic inhibition only produced behavioral impairments when rats were forced to use the context to dictate the current A-O associations. Overall, these results suggest that, while hippocampus is not required for goal-directed behavior per se, it *is* required when the context is used to guide action selection.

Our work confirms and extends previous findings showing that the context can enter into hierarchical relationships with instrumental actions (Trask and Bouton, 2014). In both the current study and that of Trask and Bouton (2014), the two contexts (Context A and Context B) differed not only in their contextual information but also in the physical actions that were used in each context. That is, four distinct actions were used during training. It is therefore possible that, instead of forming hierarchical associations whereby the context is used to decipher the ambiguous relationships between A1 and A2 and O1 and O2, rats use action-specific cues and learn two sets of independent binary associations (e.g., A1-O1, A2-O2 in Context A and A3-O1, A4-O2 in Context B). At test, these same actions were present and, thus, rats could use the action-specific cues to retrieve the binary associations learned during training.

To rule out this possibility, we also tested our rats in other cages that contained the relevant contextual information but novel levers (Experiments 2a and 2b). This meant that rats had not learned about any action-specific cues or formed specific binary associations between these levers and the outcomes they delivered during training. When tested with these novel levers, control rats were indeed able to use the context to select the correct action, suggesting that rats had formed hierarchical context-(A-O) associations and were using these associations at test to choose between the two actions. By contrast, when tested in these cages in the absence of action-specific cues, we observed deficits in outcome devaluation in rats with chemogenetic inhibition. Specifically, we found that inhibition of vHPC during training or during test rendered rats unable to bias their responding according to the context, and dHPC inhibition during test but not during training also abolished context-dependent responding. These results indicate that vHPC is required to both encode and retrieve context-(A-O) associations, while dHPC is required only for retrieval.

It should be noted that we did consistently observe an overall decrease in responding when rats were tested outside of their training cages. This decrease may be due to the context switch effect whereby a reduction in instrumental responding is observed following a change of physical context (Bouton et al., 2011, 2014; Todd, 2013; Bouton and Todd, 2014) or simply due to some extinction of the response after repeated unrewarded tests. Dorsal and ventral hippocampus have also been implicated in novelty detection and novel environment exploration (Fanselow and Dong, 2010; Duffy et al., 2013; Fredes et al., 2021; Park et al., 2021; Gómez-Ocádiz et al., 2022) and chemogenetic inhibition of either dHPC or vHPC may have impaired the rats’ ability to recognize that they were in a new cage. However, this alone cannot account for the pattern of results. First, decreased novelty detection would likely lead to less exploration and, thus, less competition with lever-press responding and, second, we observed impaired responding in rats that received DCZ during training but vehicle at test.

Importantly, chemogenetic inhibition had no effect on action selection when rats were tested in their training cages where they could potentially rely on action-specific cues and binary A-O associations (Experiments 2a and 2b), which suggests that minimizing the context-dependency of the task also renders the task hippocampus-independent. These results are also consistent with previous studies showing that inactivation or lesion of the hippocampus has no effect on goal-directed behavior, as assessed via outcome devaluation, when there is no context manipulation (Corbit et al., 2002; Macedo et al., 2008). However, interestingly, Bradfield and colleagues (2020) recently demonstrated that goal-directed actions are dependent on both the CA1 region of dHPC *and* the physical context when they are minimally trained. They observed that inhibition of dorsal CA1, during both training and test, abolished outcome devaluation after minimal, but not extended training. In addition, changing the context between training and test also led to an inability to show selective outcome devaluation for minimally trained actions. That is, dHPC is likely required for minimally trained goal-directed actions because these actions are context-dependent (Bradfield et al., 2020). It is therefore possible that, when actions are minimally trained, they enter into a hierarchical relationship with the context and, as such, both their encoding and retrieval becomes hippocampus-dependent.

Like Bradfield and colleagues (2020), we observed behavioral impairments when dHPC was inhibited during test but we didn’t observe any impairments when dHPC was inhibited during training. This may have been due to the amount of training our rats received and, if given less context-(A-O) training, it is possible that inhibition of dHPC during training *and* test would produce deficits in outcome devaluation (Bradfield et al., 2020). It also remains to be seen if the findings of Bradfield and colleagues (2020) are specific to dHPC or whether similar results might be found with vHPC inhibition. Nevertheless, taken together, these results provide compelling evidence that both dorsal and ventral hippocampus are required for goal-directed behavior but only when that behavior is context-dependent.

While both dHPC and vHPC have been implicated in the retrieval of contextual information to guide behavior (Holt and Maren, 1999; Hobin et al., 2006), one interesting aspect of the current findings is the dichotomy between these two subregions in the encoding of context-(A-O) associations. Recent attempts to characterize the difference between dHPC and vHPC have shown that neurons in the dorsal part of the hippocampus respond rapidly to a new environment and represent distinct locations and events with a precise spatial fidelity that persists across time. By contrast, neuronal ensembles in the vHPC gradually change with experience and distinguish between different behaviorally-relevant contexts (Komorowski et al., 2013). Komorowski and colleagues (2013) trained rats to discriminate two odor cues, with one cue rewarded in one context and the other rewarded in a different context. vHPC neurons initially responded with poor discrimination between both contexts but become more selective across training, suggesting a role for these neurons in discriminating how events occur in different contexts and allowing the animal to retrieve the appropriate task “rule” within each specific context. In addition, vHPC cells respond more to emotionally-charged cues than dHPC cells (Keinath et al., 2014). Together, these results suggest that dHPC represents a precise spatial environment and vHPC is required to link this environmental context with relevant behavioral events. This appears consistent with the current findings showing a role for vHPC, but not dHPC, in the formation of context-(A-O) associations.

Similarly, Biane et al. (2023) recently showed that hippocampus encodes a progressive external to internal representation of environmental events along its dorsal-ventral axis, such that dHPC represents the rich external environment and vHPC integrates this representation with behaviorally relevant information during learning. This is also consistent with recent theories on the role of vHPC in behavioral inhibition, which describes the vHPC as a context-dependent guide for appropriate behavioral strategies, suppressing or promoting a specific behavior depending on the location (Bryant and Barker, 2020). Interestingly, it has also been shown that inhibition of vHPC, but not dHPC, attenuates performance in a Pavlovian contextual biconditional discrimination task (Riaz et al., 2017) and in a context-dependent two-way signalled avoidance task (Oleksiak et al., 2021).

The vHPC also sends direct projections to the medial prefrontal cortex (Jay and Witter, 1991; Verwer et al., 1997; Ishikawa and Nakamura, 2006; Liu and Carter, 2018), a region that is heavily implicated in both goal-directed behavior (Balleine and Dickinson, 1998; Corbit and Balleine, 2003; Killcross and Coutureau, 2003; Hart and Balleine, 2016; Hart et al., 2018b, 2018a) and in the use of contextual or task-setting information to guide behavioral responding (Birrell and Brown, 2000; Miller and Cohen, 2001; Haddon and Killcross, 2006b; Marquis et al., 2007; Floresco et al., 2008; Sharpe and Killcross, 2014, 2015). Specifically, the prelimbic cortex (or Area 32) is required for encoding the A-O contingency (Corbit and Balleine, 2003; Killcross and Coutureau, 2003). Here, we show that neither dHPC or vHPC is required to encode the A-O association per se (Experiment 1b) but vHPC is required for encoding a context-(A-O) association (Experiment 2b). It is possible that, during training, the hippocampus, via its ventral part, carries contextual information to the prelimbic cortex to support the learning of behavioral responses in a specific environmental context.

An important requirement of flexible behavior is the need to select the appropriate response in different environments. Indeed, failure to use contextual information often leads to inappropriate behaviors and an inability to coordinate one’s actions to achieve a goal (Cohen and Servan-Schreiber, 1992; Haddon and Killcross, 2007; Haddon et al., 2008; Lee and Lee, 2013). The current results provide further evidence that animals use the context to guide action selection when simple binary A-O associations do not suffice (Thrailkill and Bouton, 2015). Context can therefore serve as an occasion setter that signals or modulates the specific instrumental relationships that are learned in that context (Bouton and Swartzentruber, 1986; Bouton, 1993; Myers and Gluck, 1994; Lee and Lee, 2013; Urcelay and Miller, 2014; Trask et al., 2017; Abiero and Bradfield, 2021). Here, we demonstrate a role for the hippocampus in this process. More specifically, we revealed a dissociation between ventral and dorsal hippocampus, with the former playing a seemingly more critical role in the hierarchical context modulation of goal-directed behavior than the latter. These findings not only increase our understanding of the role of hippocampus in goal-directed action but also reinforce and advance current theories on the contextual control of operant behavior (Trask et al., 2017; Abiero and Bradfield, 2021; Bouton, 2021).

## Acknowledgements

This work was supported by the French National Agency for Scientific Research (CoCoChoice ANR-19-CE37-0004-07) and a BBRF award (BBRF 27402) to S.L.P. This work was also supported by a PhD fellowship from the French Minister of Higher Education, Research and Innovation (Ministère de l’Enseignement Supérieur, de la Recherche et de l’Innovation) awarded to R.P. The authors thank Y. Salafranque for expert animal care and C. Le Moine, E Coutureau, and M. Wolff for their insightful comments on an earlier version of this manuscript.

## References

Abiero AR, Bradfield LA (2021) The contextual regulation of goal-directed actions. Current Opinion in Behavioral Sciences 41:57–62.

Armbruster BN, Li X, Pausch MH, Herlitze S, Roth BL (2007) Evolving the lock to fit the key to create a family of G protein-coupled receptors potently activated by an inert ligand. Proc Natl Acad Sci U S A 104:5163–5168.

Balleine BW, Dickinson A (1998) Goal-directed instrumental action: contingency and incentive learning and their cortical substrates. Neuropharmacology 37:407–419.

Barfield ET, Gerber KJ, Zimmermann KS, Ressler KJ, Parsons RG, Gourley SL (2017) Regulation of actions and habits by ventral hippocampal trkB and adolescent corticosteroid exposure Csicsvari J, ed. PLoS Biol 15:e2003000.

Barfield ET, Gourley SL (2019) Glucocorticoid-sensitive ventral hippocampal-orbitofrontal cortical connections support goal-directed action – Curt Richter Award Paper 2019. Psychoneuroendocrinology 110:104436.

Biane JS, Ladow MA, Stefanini F, Boddu SP, Fan A, Hassan S, Dundar N, Apodaca-Montano DL, Zhou LZ, Fayner V, Woods NI, Kheirbek MA (2023) Neural dynamics underlying associative learning in the dorsal and ventral hippocampus. Nat Neurosci:1–12.

Birrell JM, Brown VJ (2000) Medial Frontal Cortex Mediates Perceptual Attentional Set Shifting in the Rat. J Neurosci 20:4320–4324.

Bouton ME (1993) Context, time, and memory retrieval in the interference paradigms of Pavlovian learning. Psychol Bull 114:80–99.

Bouton ME (2021) Context, attention, and the switch between habit and goal-direction in behavior. Learn Behav 49:349–362.

Bouton ME, Swartzentruber D (1986) Analysis of the associative and occasion-setting properties of contexts participating in a Pavlovian discrimination. Journal of Experimental Psychology: Animal Behavior Processes 12:333–350.

Bouton ME, Todd TP (2014) A fundamental role for context in instrumental learning and extinction. Behav Processes 104:13–19.

Bouton ME, Todd TP, Leon SP (2014) Contextual control of discriminated operant behavior. Journal of experimental psychology Animal learning and cognition 40:92–105.

Bouton ME, Todd TP, Vurbic D, Winterbauer NE (2011) Renewal after the extinction of free operant behavior. Learn Behav 39:57–67.

Bradfield LA, Balleine BW (2013) Hierarchical and binary associations compete for behavioral control during instrumental biconditional discrimination. Journal of Experimental Psychology: Animal Behavior Processes 39:2–13.

Bradfield LA, Leung BK, Boldt S, Liang S, Balleine BW (2020) Goal-directed actions transiently depend on dorsal hippocampus. Nat Neurosci Available at: http://www.nature.com/articles/s41593-020-0693-8 [Accessed August 20, 2020].

Bryant KG, Barker JM (2020) Arbitration of Approach-Avoidance Conflict by Ventral Hippocampus. Front Neurosci 14:615337.

Cerpa JC, Piccin A, Dehove M, Lavigne M, Kremer EJ, Wolff M, Parkes SL, Coutureau E (2023) Inhibition of noradrenergic signalling in rodent orbitofrontal cortex impairs the updating of goal-directed actions Bradfield LA, Wassum KM, Bradfield LA, eds. eLife 12:e81623.

Cohen JD, Servan-Schreiber D (1992) Context, cortex, and dopamine: a connectionist approach to behavior and biology in schizophrenia. Psychol Rev 99:45–77.

Colwill RM, Rescorla RA (1990) Evidence for the hierarchical structure of instrumental learning. Animal Learning & Behavior 18:71–82.

Corbit LH, Balleine BW (2003) The role of prelimbic cortex in instrumental conditioning. Behav Brain Res 146:145–157.

Corbit LH, Ostlund SB, Balleine BW (2002) Sensitivity to instrumental contingency degradation is mediated by the entorhinal cortex and its efferents via the dorsal hippocampus. J Neurosci 22:10976–10984.

Duffy AM, Schaner MJ, Chin J, Scharfman HE (2013) Expression of c-fos in hilar mossy cells of the dentate gyrus in vivo. Hippocampus 23:649–655.

Fanselow MS, Dong H-W (2010) Are the Dorsal and Ventral Hippocampus Functionally Distinct Structures? Neuron 65:7–19.

Floresco SB, Block AE, Tse MTL (2008) Inactivation of the medial prefrontal cortex of the rat impairs strategy set-shifting, but not reversal learning, using a novel, automated procedure. Behavioural Brain Research 190:85–96.

Fredes F, Silva MA, Koppensteiner P, Kobayashi K, Joesch M, Shigemoto R (2021) Ventro-dorsal Hippocampal Pathway Gates Novelty-Induced Contextual Memory Formation. Current Biology 31:25–38.e5.

Gómez-Ocádiz R, Trippa M, Zhang C-L, Posani L, Cocco S, Monasson R, Schmidt-Hieber C (2022) A synaptic signal for novelty processing in the hippocampus. Nat Commun 13:4122.

Gourley SL, Lee AS, Howell JL, Pittenger C, Taylor JR (2010) Dissociable regulation of instrumental action within mouse prefrontal cortex. Eur J Neurosci 32:1726–1734.

Haddon JE, George DN, Killcross S (2008) Contextual control of biconditional task performance: Evidence for cue and response competition in rats. Quarterly Journal of Experimental Psychology 61:1307– 1320.

Haddon JE, Killcross S (2006a) Prefrontal cortex lesions disrupt the contextual control of response conflict. J Neurosci 26:2933–2940.

Haddon JE, Killcross S (2006b) Prefrontal Cortex Lesions Disrupt the Contextual Control of Response Conflict. J Neurosci 26:2933–2940.

Haddon JE, Killcross S (2007) Contextual Control of Choice Performance. Annals of the New York Academy of Sciences 1104:250–269.

Hart G, Balleine BW (2016) Consolidation of Goal-Directed Action Depends on MAPK/ERK Signaling in Rodent Prelimbic Cortex. The Journal of neuroscience: the official journal of the Society for Neuroscience 36:11974–11986.

Hart G, Bradfield LA, Balleine BW (2018a) Prefrontal Corticostriatal Disconnection Blocks the Acquisition of Goal-Directed Action. The Journal of neuroscience: the official journal of the Society for Neuroscience 38:1311–1322.

Hart G, Bradfield LA, Fok SY, Chieng B, Balleine BW (2018b) The Bilateral Prefronto-striatal Pathway Is Necessary for Learning New Goal-Directed Actions. Current biology: CB 28:2218–2229 e7.

Ishikawa A, Nakamura S (2006) Ventral Hippocampal Neurons Project Axons Simultaneously to the Medial Prefrontal Cortex and Amygdala in the Rat. Journal of Neurophysiology 96:2134–2138.

Jay TM, Witter MP (1991) Distribution of hippocampal CA1 and subicular efferents in the prefrontal cortex of the rat studied by means of anterograde transport of Phaseolus vulgaris-leucoagglutinin. J Comp Neurol 313:574–586.

Keinath AT, Wang ME, Wann EG, Yuan RK, Dudman JT, Muzzio IA (2014) Precise spatial coding is preserved along the longitudinal hippocampal axis. Hippocampus 24:1533–1548.

Killcross S, Coutureau E (2003) Coordination of actions and habits in the medial prefrontal cortex of rats. Cerebral cortex 13:400–408.

Komorowski RW, Garcia CG, Wilson A, Hattori S, Howard MW, Eichenbaum H (2013) Ventral Hippocampal Neurons Are Shaped by Experience to Represent Behaviorally Relevant Contexts. Journal of Neuroscience 33:8079–8087.

Lee I, Lee CH (2013) Contextual behavior and neural circuits. Front Neural Circuits 7:84.

Liu X, Carter AG (2018) Ventral Hippocampal Inputs Preferentially Drive Corticocortical Neurons in the Infralimbic Prefrontal Cortex. J Neurosci 38:7351–7363.

Macedo CE, Sandner G, Angst M-J, Guiberteau T (2008) Rewarded associative and instrumental conditioning after neonatal ventral hippocampus lesions in rats. Brain Research 1215:190–199.

Maren S, Phan KL, Liberzon I (2013) The contextual brain: implications for fear conditioning, extinction and psychopathology. Nat Rev Neurosci 14:417–428.

Marquis J-P, Killcross S, Haddon JE (2007) Inactivation of the prelimbic, but not infralimbic, prefrontal cortex impairs the contextual control of response conflict in rats. Eur J Neurosci 25:559–566.

Miller EK, Cohen JD (2001) An integrative theory of prefrontal cortex function. Annu Rev Neurosci 24:167– 202.

Myers CE, Gluck MA (1994) Context, conditioning, and hippocampal rerepresentation in animal learning. Behav Neurosci 108:835–847.

Nagai Y et al. (2020) Deschloroclozapine, a potent and selective chemogenetic actuator enables rapid neuronal and behavioral modulations in mice and monkeys. Nat Neurosci 23:1157–1167.

Nentwig TB, Obray JD, Vaughan DT, Chandler LJ (2021) Behavioral and slice electrophysiological assessment of DREADD ligand, deschloroclozapine (DCZ) in rats.:2021.10.25.465454 Available at: https://www.biorxiv.org/content/10.1101/2021.10.25.465454v1 [Accessed March 11, 2022].

Oleksiak CR, Ramanathan KR, Miles OW, Perry SJ, Maren S, Moscarello JM (2021) Ventral hippocampus mediates the context-dependence of two-way signaled avoidance in male rats. Neurobiol Learn Mem 183:107458.

Park AJ, Harris AZ, Martyniuk KM, Chang C-Y, Abbas AI, Lowes DC, Kellendonk C, Gogos JA, Gordon JA (2021) Reset of hippocampal-prefrontal circuitry facilitates learning. Nature 591:615–619.

Park CHJ, Ganella DE, Perry CJ, Kim JH (2020) Dissociated roles of dorsal and ventral hippocampus in recall and extinction of conditioned fear in male and female juvenile rats. Exp Neurol 329:113306.

Pinizzotto CC, Heroux NA, Horgan CJ, Stanton ME (2020) Role of dorsal and ventral hippocampal muscarinic receptor activity in acquisition and retention of contextual fear conditioning. Behav Neurosci 134:460–470.

Rescorla RA (1992) Hierarchical Associative Relations in Paviovian Conditioning and Instrumental Training.

Riaz S, Schumacher A, Sivagurunathan S, Van Der Meer M, Ito R (2017) Ventral, but not dorsal, hippocampus inactivation impairs reward memory expression and retrieval in contexts defined by proximal cues: RIAZ et al. Hippocampus 27:822–836.

Rogan SC, Roth BL (2011) Remote control of neuronal signaling. Pharmacol Rev 63:291–315.

Rudy JW (2009) Context representations, context functions, and the parahippocampal-hippocampal system. Learning & Memory 16:573–585.

Sharpe M, Killcross S (2015) The prelimbic cortex uses contextual cues to modulate responding towards predictive stimuli during fear renewal. Neurobiol Learn Mem 118:20–29.

Sharpe MJ, Killcross S (2014) The Prelimbic Cortex Contributes to the Down-Regulation of Attention Toward Redundant Cues. Cerebral Cortex 24:1066–1074.

Thrailkill EA, Bouton ME (2015) Contextual control of instrumental actions and habits. Journal of experimental psychology Animal learning and cognition 41:69–80.

Todd TP (2013) Mechanisms of renewal after the extinction of instrumental behavior. Journal of Experimental Psychology: Animal Behavior Processes 39:193–207.

Trask S, Bouton ME (2014) Contextual control of operant behavior: evidence for hierarchical associations in instrumental learning. Learning & behavior 42:281–288.

Trask S, Thrailkill EA, Bouton ME (2017) Occasion setting, inhibition, and the contextual control of extinction in Pavlovian and instrumental (operant) learning. Behavioural Processes 137:64–72.

Urcelay GP, Miller RR (2014) The functions of contexts in associative learning. Behav Processes 0:2–12.

Verwer RW, Meijer RJ, Van Uum HF, Witter MP (1997) Collateral Projections from the Rat Hippocampal Formation to the Lateral and Medial Prefrontal Cortex. Hippocampus 7:397–402.

Wang ME, Fraize NP, Yin L, Yuan RK, Petsagourakis D, Wann EG, Muzzio IA (2013) Differential roles of the dorsal and ventral hippocampus in predator odor contextual fear conditioning. Hippocampus 23:451– 466.

Yoshida K, Drew MR, Kono A, Mimura M, Takata N, Tanaka KF (2021) Chronic social defeat stress impairs goal-directed behavior through dysregulation of ventral hippocampal activity in male mice. Neuropsychopharmacology 46:1606–1616.

Yoshida K, Drew MR, Mimura M, Tanaka KF (2019) Serotonin-mediated inhibition of ventral hippocampus is required for sustained goal-directed behavior. Nat Neurosci 22:770–777.

